# Cortical network fingerprints predict deep brain stimulation outcome in dystonia

**DOI:** 10.1101/470526

**Authors:** Gabriel Gonzalez-Escamilla, Muthuraman Muthuraman, Martin M Reich, Nabin Koirala, Christian Riedel, Martin Glaser, Florian Lange, Günther Deuschl, Jens Volkmann, Sergiu Groppa

## Abstract

**Background:** Deep brain stimulation (DBS) is an effective evidence-based therapy for dystonia. However, no unequivocal predictors of therapy responses exist. We investigate whether patients optimally responding to DBS present distinct brain network organization and structural patterns.

**Methods:** Based on a German multicentre cohort of eighty-two dystonia patients with segmental and generalized dystonia, who received DBS implantation in the globus pallidus internus patients were classified based on the clinical response 36 months after DBS, as superior-outcome group or moderate-outcome group, as above or below 70% motor improvement, respectively. Fifty-one patients met MRI-quality and treatment response requirements (mean age 51.3 ± 13.2 years; 25 female) and were included into further analysis. From preoperative MRI we assessed cortical thickness and structural covariance, which were then fed into network analysis using graph theory. We designed a support vector machine to classify subjects for the clinical response based on group network properties and individual grey matter fingerprints.

**Results:** The moderate-outcome group showed cortical atrophy mainly in the sensorimotor and visuomotor areas and disturbed network topology in these regions. From all the structural integrity of the cortical mantle explained about 45% of the stimulation amplitude. Classification analyses achieved 88% of accuracy using individual grey matter atrophy patterns to predict DBS outcome.

**Conclusions:** The analysis of cortical integrity and network properties could be developed into independent predictors to identify dystonia patients who benefit from DBS.

## Introduction

Deep brain stimulation (DBS) is a well-established treatment for patients with medically intractable, segmental and generalized dystonia, for which the globus pallidus internus (GPi-DBS) is an efficient target,[1 2]. However, the degree of improvement varies among patients. Here, we postulate that structural brain network properties as derived from MRI may act as predictors in dystonia patients undergoing DBS. Furthermore, elucidating the neuroanatomical basis for network dysfunction in dystonia would have a direct implication for the surgical intervention,[3–5] and is a critical first step towards developing personalized therapeutic solutions and effective neuromodulation paradigms. An individualized characterization of abnormal anatomical and physiological networks in each patient could lead to risk minimization for those patients who might be susceptible to poor DBS outcome due to specific disease fingerprints or irreversible secondary abnormalities in the brain circuits or periphery.

Brain circuit alterations have been attested in patients with dystonia in several brain regions,[6 7], leading to the notion that the disease cannot arise from damage of a single structure, but rather from a network dysfunction,[8 9]. This network dysfunction leads to excessive movement that is normalized under DBS,[10]. Here, we investigate how preoperative brain network properties relate to the clinical outcome of GPi-DBS. For this, we reconstruct grey matter cerebral networks using graph theory to quantify local and global structural fingerprints in patients with segmental and generalized dystonia.

Graph theory has become a relevant tool to explore brain circuit abnormalities in neuropsychiatric disorders and to quantify patterns of disease-related reorganization,[11 12]. Small-world properties, which have been related to physiological brain functioning and reflect a clustered network with short paths, offer a basis for maintained network functionality and efficiency,[13 14]. Despite the growing interest in DBS, still much remains unknown about the network-level impact of neuromodulatory effects. In this study, we sought to identify structural fingerprints that predict the response and maintenance of benefit in a stable clinical state after 3 years of GPi-DBS using a novel computational approach consisting of graph theory and machine learning techniques.

## Methods

### Standard protocol approvals, registrations, and patient consents

This retrospective study was approved by the institutional review and Ethics boards at each participating centre and was carried out in accordance with the Declaration of Helsinki. All patients provided written informed consent.

### Study participants and data acquisition

For a multicentric study including 82 primary dystonia patients who underwent GPi-DBS, two separate group of dystonia patients were analysed according to the recruitment and MRI image quality (see the results section below). This included a main cohort of patients treated at the University Clinic of Kiel (N = 36, mean age ± SD = 50.1 ± 12.1 years, 16 females) and a replication group from the University Clinic Würzburg (N = 15, mean age ± SD = 54.2 ± 9.3 years, 9 females). None of the patients presented secondary dystonia. The precise description can be found elsewhere,[1 2].

For the main dystonia cohort, patients underwent a T1-weighted magnetization-prepared rapid gradient-echo (MP-RAGE) MRI (repetition time (TR) = 10.83 ms; echo time (TE) = 2.02 ms; flip angle = 30°; 2.0 mm effective slice thickness; acquisition matrix = 256 x 256) using a 1.5T Philips Achieva scanner with an 8-channel SENSE head coil. For the replication cohort, again, high resolution T1-weighted MP-RAGE images were acquired (TR = 11 ms, TE = 4.92 ms, flip angle = 20°, slice thickness = 1mm, Acquisition matrix = 256 × 256) with a 3T Siemens TrioTim scanner, using a 32-channel SENSE head coil.

### GPi-DBS electrode implantation and clinical outcomes

All patients were implanted with bilateral electrodes (model 3387, Medtronic) into the posterior-ventral portion of the internal globus pallidus. The exact neurosurgical procedure is described elsewhere.[1 15] Standard stereotactic coordinates for anatomical targeting were individually adapted by direct visualization of the GPi on the MR images. Stimulation parameters including amplitude, frequency, and pulse width were adjusted for each individual patient. The effects of DBS on clinical outcomes were quantified as the improvement percentage in the movement scale of the Burke–Fahn–Marsden Dystonia Rating Scale (BFMDRS) for generalized dystonia,[2] and the Toronto Western Spasmodic Torticollis Rating Scale (TWSTRS) for torticollis,[1] assessed before and three years after surgery. The improvement percentage at follow-up was further used to classify the patients as superior-outcome group (SOG) and moderate-outcome group (MOG). The stimulation adjustment and the clinical evaluation were performed by clinicians who were blinded to the hypothesis and goals of this study.

### Cortical thickness maps and structural connectivity measures

All T1 images were pre-processed using the automatic surface-based pipeline of FreeSurfer (v5.3, http://surfer.nmr.mgh.harvard.edu), which included: skull stripping, image affine registration, bias correction and segmentation of grey and white matter tissue compartments, separation of brain hemispheres and subcortical structures, and construction of smooth representation of the grey/white interface and the cortical surface,[16]. The distance between the corresponding vertices of inner and outer cortical surfaces was then used as a measure of cortical thickness at each vertex.

The spatial correlation of cortical thickness for each pair of the 68 regions in the Desikan-Killiany atlas[17] was compiled to form a morphometric similarity matrix and analysed using graph theory. The covariance matrix of each group was then binarised with a network-derived threshold, where an entry is 1 if the correlation weighting between a pair of regions is greater than a minimum density threshold i.e. the density at which all the regions are fully connected in the network of each group. This ensures that none of the networks are fragmented. Consistent with previous studies,[18], the association matrices were thresholded at a range of network densities from the minimum density in steps of .5% across a 10% degree range. This was done to ensure that group differences are not confounded by differing number of nodes and edges due to an absolute threshold at a single density. Following, to evaluate network topology, the small-world index was calculated based on two key measures: The clustering coefficient (C), a global count of the interconnectedness (number of connections) between the network regions and their nearest neighbours; and the characteristic path length (P_L_), which is the average minimum distance to connect all pairs of regions in the network. P_L_ and C were further normalized in comparison to a random network to avoid the influence of other topological characteristics, leading to the respective parameters “lambda” and “gamma”. The network small-worldness, “sigma”, was then calculated as the ratio of gamma to lambda (sigma = lamdda/gamma),[19]. Additionally, the network’s local efficiency (E_local_), the nodal degree centrality (count of how many neighbours a single region has) and nodal clustering (clustering coefficient of every single region) were used to evaluate local connectivity differences. See [19] for detailed mathematical definitions.

### Statistical analyses

Differences in cortical thickness between the two groups (SOG vs MOG) as well as associations between cortical thickness and stimulation parameters were statistically determined using the general linear model (GLM) with age and gender as nuisance variables using a threshold of *p* < 0.05 corrected for multiple comparisons with 10,000 Monte Carlo Z-simulations. Similarly, the volume of subcortical structures was compared and corrected with FDR (*p* < 0.05).

The between-group differences in each network topology measure (C, P_L_, sigma, E_local_), were tested via *t*-test at p < 0.05. Further, a non-probabilistic binary support vector machine (SVM) classifier was used to test whether a particular network parameter or regional cortical thickness measure would better predict the DBS outcome. Briefly, the SVM algorithm looks for an optimally separating threshold between the two data sets by maximizing the margin between the classes’ closest points. The points lying on the boundaries are called support vectors, and the middle of the margin is the optimal separating threshold. Here, we have used the polynomial function kernel for this projection due to its good performance,[20]. The SVM selection was performed using 75% of the sample as training, and a 10-fold cross-validation to compute the correct classification ration (CCR). Further, the area under the curve (AUC) from Receiver Operating Characteristic (ROC) curves was adopted to ensure the sensitivity of the network measures and measures of cortical thickness as predictors of GPi-DBS responsiveness in the dystonia patients.

All analyses between the two groups of interest (SOG vs MOG) were conducted independently in the two study cohorts. Therefore, figures and tables summarize the results in the main population, and that were replicated aftermath in the second cohort.

### Data availability

The datasets generated during and/or analysed during the current study will be made available from the corresponding author on reasonable request.

## Results

### Demographics

From the initial population (N = 82) of this multicentric study, 31 patients were not eligible due to differences in the MR image quality. The remaining 51 patients were then classified upon their sustained clinical improvement three years after GPi-DBS, into two demographically equivalent groups of SOG (N = 26, 19 in the main cohort and 7 in the replication group) or MOG (N = 25; 17 and 8, from each cohort respectively), see Table 1 for more details on the group distributions and demographics.

**Table 1.** Group demographics.

|  | Main sample (1.5T MRI) |  | Replication (3T MRI) |  |
| --- | --- | --- | --- | --- |
|  | moderate-outcome<br>group<br>(N=17) | superior-outcome<br>group<br>(N=19) | moderate-outcome<br>group<br>(N=8) | superior-outcome<br>group<br>(N=7) |
| **Generalized/cervical dystonia** | 9/8 | 10/9 | 1/7 | 5/2 |
| **Motor score (SD) – pre-DBS** | 34.4 (15.9) | 41.1 (29.5) | 23.13 (3.9) | 30.2 (11.4) |
| **Motor score (SD) – follow-up** | 21.2 (17.8) | 5.3 (5.7) | 15.9 (6.3) | 6.3 (4) |
| **Motor improvement (%) (SD)** | 40.4 (26.3) | 85.8 (8.6) | 30.8 (21.2) | 80.2 (9.5) |
| **female/male** | 7/10 | 9/10 | 5/3 | 4/3 |
| **Age (SD)** | 52.8 (11.4) | 47.7 (12.3) | 62.1 (9.3) | 45.1 (17.4) |

SD, standard deviation of the mean

### Preoperative anatomical and network fingerprints

Correspondence of between-group differences in cortical thickness among populations (Fig 1) was shown for cortical thinning in the superior, middle and inferior (ventral motor area) frontal, paracentral and parietal (precuneus) regions in MOG. Accordingly, the regression analyses showed a negative association between the cortical thickness of these areas and the DBS stimulation amplitude for the left (r = 0.46, p = 0.006) and the right (r = 0.44, p = 0.007) hemispheres. Subcortically, MOG also showed reduced volumes of several basal ganglia system structures and the thalamus (Table 2).

**Figure 1.**
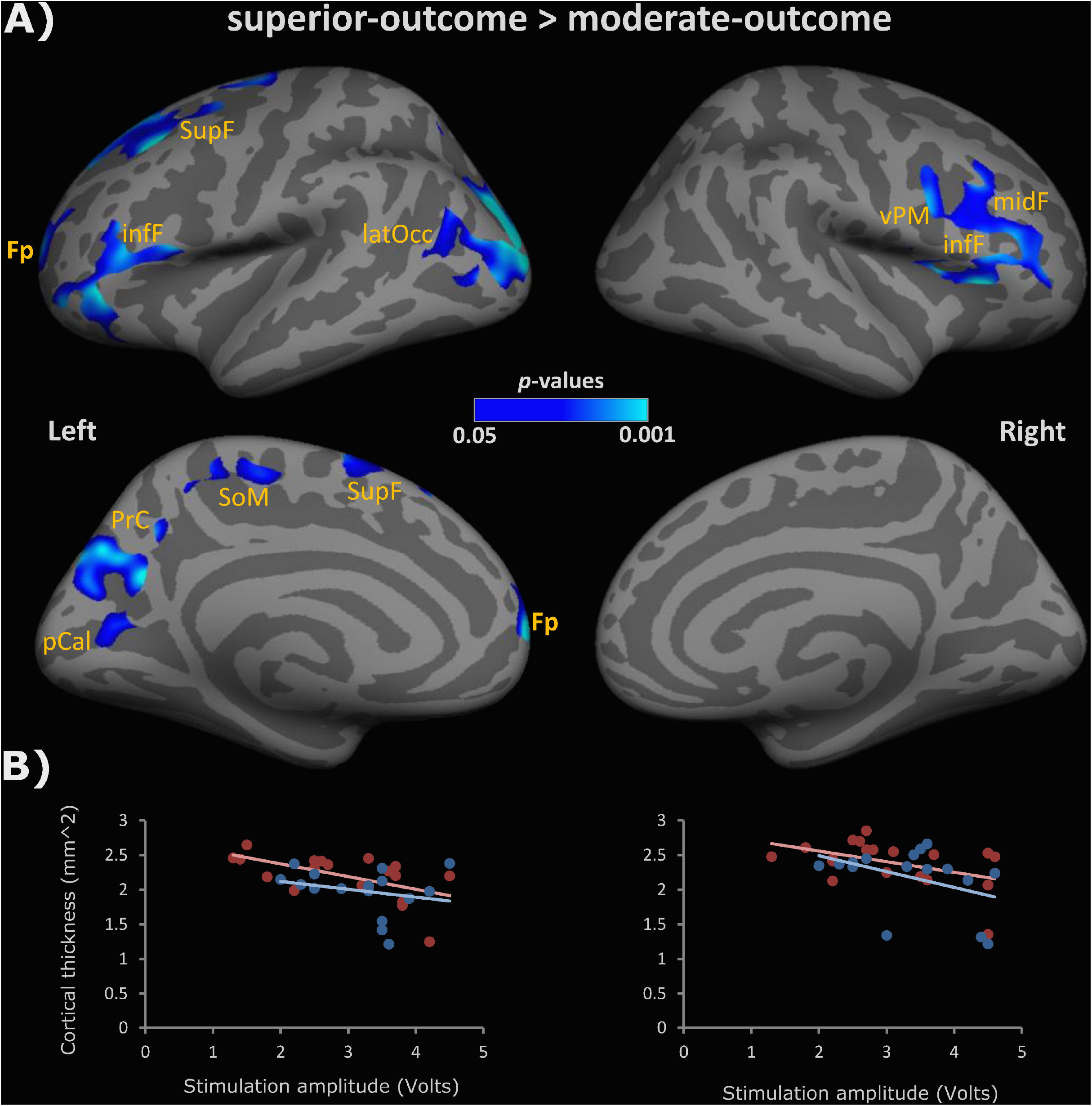
Cortical thinning in the brain of the moderate-outcome group. **A:** Overlapping map showing the regions where GPi-DBS moderate-outcome dystonia patients showed significantly decreased cortical thickness when compared to GPi-DBS the superior-outcome group. Top row lateral hemisphere surfaces, bottom row medial hemisphere surfaces. Fp, frontal pole; infF, inferior frontal; SupF, superior frontal; latOcc, lateral occipital; SoM, sensorimotor area (paracentral); PrC, precuneus; pCal, pericalcarine; vPM, ventral primary motor; midF, middle frontal. **B:** Regression plots showing the negative association between cortical thickness and the amplitude of the DBS stimulation for the superior-outcome group (red) and the moderate-outcome group (blue). Nuisance variables: age and gender; p-values are corrected for multiple comparisons using Monte Carlo Z-simulations at *p* < 0.05.

**Table 2.** Group differences in GM subcortical volumes between GPi-DBS superior-outcome and moderate-outcome groups.

| GM nuclei | Volume (mm<sup>3</sup>) |  | p value | t stat |
| --- | --- | --- | --- | --- |
|  | moderate-outcome group | superior-outcome group |  |  |
| lh Thalamus | 8112.75 | 8698.57 | **0.036** | -1.84 |
| lh Caudate | 2994.93 | 3320.32 | **0.012** | -2.32 |
| lh Putamen | 4041.22 | 4470.00 | **0.036** | -1.84 |
| lh Pallidum | 1370.11 | 1486.23 | **0.049** | -1.69 |
| lh Hippocampus | 3586.46 | 3700.96 | 0.26 | -0.64 |
| lh Amygdala | 1302.18 | 1353.27 | 0.21 | -0.83 |
| lh Cerebellum ctx | 47547.32 | 49629.38 | 0.16 | -0.99 |
| rh Thalamus | 7458.66 | 7887.13 | **0.042** | -1.76 |
| rh Caudate | 2697.32 | 3050.44 | **0.043** | -1.76 |
| rh Putamen | 3919.97 | 4273.07 | **0.029** | -1.94 |
| rh Pallidum | 1350.42 | 1446.20 | **0.048** | -1.69 |
| rh Hippocampus | 3689.66 | 4155.62 | **0.022** | -2.08 |
| rh Amygdala | 1371.93 | 1389.26 | 0.38 | -0.30 |
| rh Cerebellum ctx | 48459.98 | 49288.93 | 0.36 | -0.37 |

lh, left hemisphere; rh, right hemisphere; ctx, cortex.

Compared with SOG, MOG showed decreased small-worldness (sigma, t = 4.7, *p* = 1.8e-5) and lambda (t = 3.82, *p* = 0.0002), and increased gamma (t = 3.81, *p* = 0.005) and E_local_ (t = 1.6, *p* = 0.05), indicating long-range disconnection and higher local connectivity in clustered, neighbouring areas (Fig 2A). The regional analyses revealed increased degree centrality in the central and fronto-parietal regions in MOG (Fig 2B). Group differences in regional clustering coefficient were also observed in central and fronto-parietal regions (Fig 2C).

**Figure 2.**
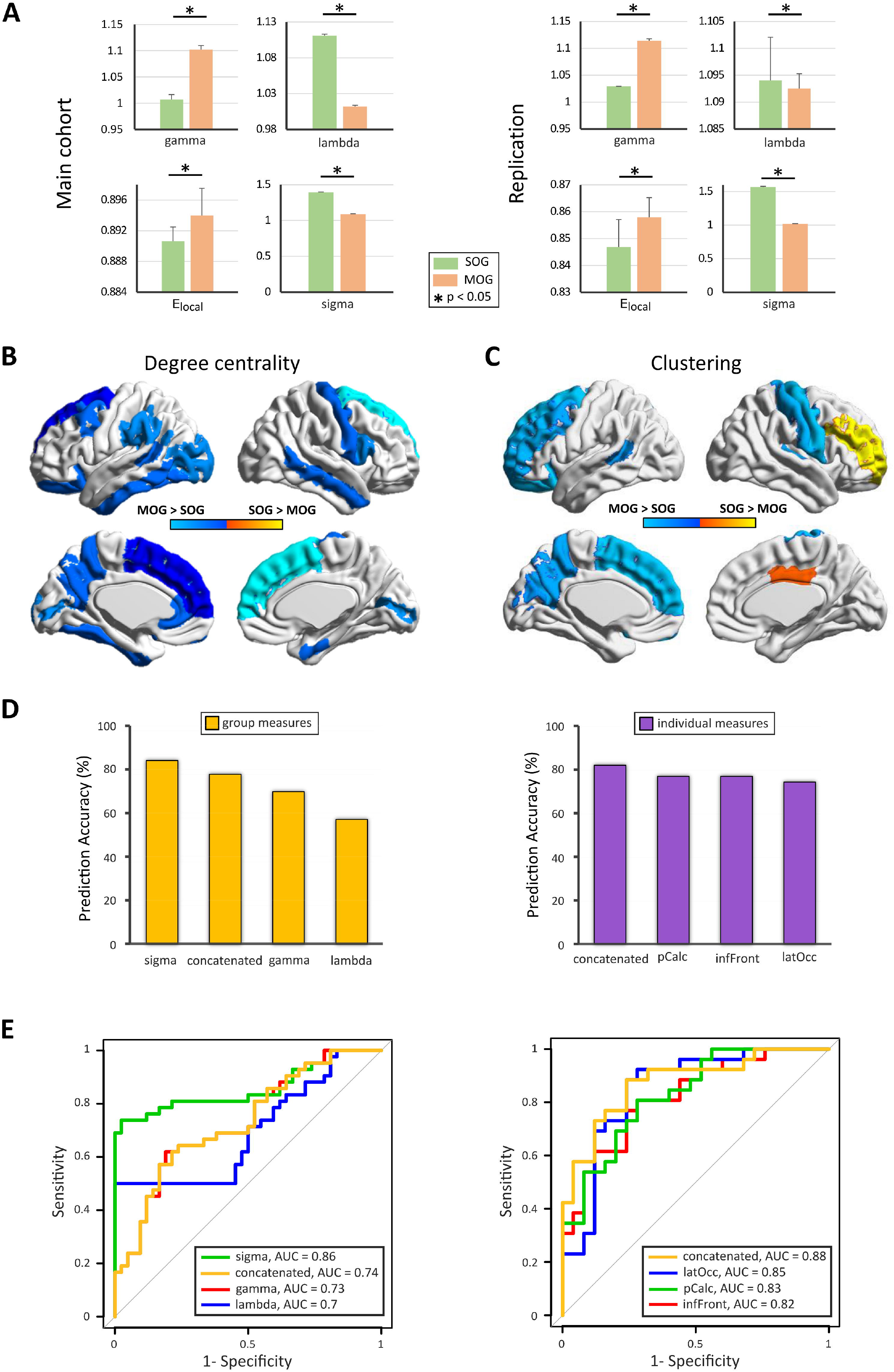
Network and prediction analyses. **A:** Between-group differences in network parameters for the main (left column), and replication (right column) cohorts. **B:** Most important regions for structural topology as determined by degree centrality. **C:** Regional differences in clustering coefficient, cold colours indicate regions with higher clustering coefficient in MOG than in SOG, warm colours indicate the opposite. **D:** Prediction accuracy of the support vector machine at training, test and overall for gamma, lambda and sigma parameters as well as the combined accuracy. **E:** Receiver Operating Characteristic (ROC) curves and the respective area under the curve (AUC) for gamma, lambda and sigma (left side) and individual regional integrity (right side). latOcc: lateral occipital; pCalc: pericalcarine; inFront: inferior frontal; MOG: moderate-outcome group; SOG: superior-outcome group.

### Predictors of GPi-DBS responsiveness

Careful review of postoperative images showing the electrode location did not revealed any obvious deviation of the target in SOG and MOG patients.

The designed group prediction model using the network measures to stratify the patients for their clinical outcome (Fig 2D and E) showed the highest test accuracy for sigma (CCR = 84.1%; AUC = 0.86), followed by lambda (CCR = 69.8%; AUC = 0.7) and gamma (CCR = 57.1%; AUC = 0.73) with a mean accuracy (CCR = 77.8%; AUC = 0.73) for the concatenated variable.

For the individual measures the three regions with the highest accuracy were: left inferiorfrontal (CCR = 74.4%; AUC = 0.82), left lateraloccipital (CCR = 76.9%; AUC = 0.85) and left pericalcarine (CCR = 76.9%; AUC = 0.83) with the best accuracy for the concatenated variable (CCR = 82.1%; AUC = 0.88), demonstrating that network topology and cortical atrophy have substantial influence on clinical outcome.

## Discussion

Grey matter network properties predict the clinical outcome to GPi-DBS in patients with dystonia. SOG presented brain circuits with a small-world topology. Patients with atrophy in a wide-spread network of association, sensorimotor and visuomotor areas have disturbed network architecture and a worse long-term outcome. Furthermore, increased local connectivity was associated with worse clinical outcome.

In the current study the classification analyses allowed a very robust delimitation of superior-outcome versus moderate-outcome groups both at the group and single subject level. Network properties and regional grey matter integrity information therefore represent putative correlates specific to brain behavior in patients with dystonia and is related to functional neuromodulation,[10]. These network abnormalities are likely caused by a reorganization of the parietal to frontal connections, with modified cortico-cortical or cortico-subcortical visuomotor and sensory information processing. These have been repeatedly related to abnormal motor control and generation of dystonic movements,[21]. This was further evidenced by the morphometric alterations in the basal ganglia and thalamus in MOG.

Alterations within the sensorimotor and associative circuits, involving central, frontal, and parietal cortices have been reported, suggesting that dystonia may represent a disorder of large-scale networks as opposed to basal ganglia pathology alone,[6 8]. In support of this novel view, a loss of long range connections was shown in MOG to DBS. Furthermore, in the MOG, a regional increase in degree of centrality and clustering coefficient, interpreted as an increased susceptibility of these particular regions to cause network failures, corresponded with regional structural alterations, likely reducing the systemic neuromodulatory effectiveness of GPi-DBS. This further implies that the optimal trade-off between wiring-cost minimization and efficiency of information transfer may play a key role in the outcome of GPi-DBS interventions in dystonia patients.

Computational studies have demonstrated that small-world network architecture requires specific control strategies allowing the enhancement of recovery following system perturbations,[22]. Here, we show that motor, sensorimotor and associative regions with impaired microstructure and connectivity (i.e. increased clustering) disturb the physiological motor control and counteract the normalization of dystonic movements to DBS.[13] Thus, the neuromodulation provided by DBS stimulation exerts specific effects on ongoing brain networks activity[23] and its efficacy depends not only on the local stimulation target, but relies moreover on the network characteristics.

A limitation of our study could be the fact that the electrode localization is an important confounder for DBS outcomes[24] in our patients the position of the implant electrodes was projected to the preoperative images and visually checked retrospectively. No clinically relevant shifts of the target that have made a reimplantation needed have been noticed in SOG and MOG. Additionally, the DYT-1 status has also been reported to be associated with treatment outcome[25] this distinction that was not made in our patients. Despite previous studies have shown that both, dystonia patients with DYT-1 and without a known genetic cause showed marked improvement after GPi-DBS,[26]. Future studies should control for this in larger populations.

Overall, our study shows the strong emerging potential of structural network studies in predicting GPi-DBS outcomes at the group and single subject level, which in turn can be used for personalized therapeutic approaches when selecting patients who are likely to benefit from this therapy.

Author Contributions
G.G.E., M.M., and S.G. designed the study, validated the extracted data, supervised the study, made a critical revision of the manuscript for intellectual content, and approved the final version of the manuscript. M.M.R., J.K., and S.G. acquired and extracted the data. G.G.E., M.M., and N.K. extracted the data, and performed the MRI-based volumetric analysis. G.G.E. performed the statistical analysis, and drafted the manuscript for intellectual content. S.G. and M.M. made a critical revision of the manuscript for intellectual content and results interpretation. S.G. had full access to all of the data in the study and take responsibility for the integrity of the data and the accuracy of the data analysis. All authors contributed to the writing of the manuscript for intellectual content, revised and approved the final version of the manuscript.

## Acknowledgments

The authors would like to thank Cheryl Ernest for proofreading the manuscript.

